# Emergent Simplicities in Stochastic Intergenerational Homeostasis

**DOI:** 10.1101/2023.01.18.524627

**Authors:** Kunaal Joshi, Charles S. Wright, Karl F. Ziegler, Elizabeth M. Spiers, Jacob T. Crosser, Samuel Eschker, Rudro R. Biswas, Srividya Iyer-Biswas

## Abstract

How do complex systems maintain key emergent “state variables” at desired target values to within specified tolerances? This question was first posed in the context of homeostasis in living systems over a century ago, and yet the precise quantitative rules governing this phenomenon have remained fiercely debated. We herein present a direct solution through a synthesis of high-precision experiments and first principles-based physics theory. After introducing a general approach that incorporates the inherently stochastic and dynamic nature of organismal homeostasis, we provide direct experimental evidence that stochastic intergenerational homeostasis is indeed maintained. Next, we identify a series of emergent simplicities hidden in these data. Remarkably, the dynamics of intergenerational homeostasis of organismal sizes are Markovian, or history-independent. The precision data reveal an intergenerational scaling law that fully determines, with no fine-tuning parameters, the exact stochastic map governing homeostasis, as borne out by compelling data– theory matches. These emergent simplicities in turn yield the necessary and sufficient condition for stochastic homeostasis, with surprising implications for the architecture of the underlying control system. Validation across different growth conditions, cell morphologies, experimental modalities, and organisms comprehensively establishes the universality of the results presented here.

---

Homeostasis describes a self-regulating process wherein a complex system maintains internal stability despite unavoidable fluctuations in its constitutive dynamics. This concept was instrumental to the development of the field of physiology (*1*); as formulated by Bernard, stability of an organism’s “internal milieu” is not simply a feature, but indeed a prerequisite for life (*2*). Early conceptualizations philosophically permitted the possibility of stochasticity in the homeostatic variable (*2–4*). However, deterministic frameworks have since become pervasive, with organisms thought—in analogy to mechanical self-regulating apparatuses—to defend the setpoint values of homeostatic variables (*5*) against intrinsic and extrinsic fluctuations. This approach can be traced to early developments in control theory as applied to engineered inanimate systems (*6*). Within this perspective, deviations from desired setpoints are taken to indicate organismal malfunction, rather than natural variability resulting from inherent stochasticity in the dynamics (*7–12*).

Realistically, biological systems feature layered architectures and hierarchical organization, which must somehow conspire to ensure homeostasis of emergent organismal “state variables” despite the inherently probabilistic nature of the underlying biophysical dynamics (*1*). Embracing this systems design point of view, we conceptualize homeostasis as a dynamical—albeit stochastic—process, rather than as an end state (i.e., a setpoint). We further define the problem of stochastic homeostasis by distinguishing two potential routes to achieving it. We term the first option “elastic adaptation”, a passive mechanism that involves reflexive responses to instantaneous conditions with no retention of long-term memory (as in Ashby’s Homeostat (*13*), the first technical model of homeostasis). In contrast, active, “plastic adaptation” is achieved through reflective responses involving strategic integration over a finite memory of the past (as in Shannon’s Maze Machine (*14*)). Which scheme(s) do organisms use?

To identify the quantitative principles that ensure the robust and dynamical control of stochastic yet homeostatic organismal state variables, we turn to one of the simplest organisms—a bacterial cell—and one of its best-studied homeostatic state variables—cell size. Growth and division of a single bacterial cell is an inherently stochastic process with significant fluctuations (Fig. 1). Yet, current models of how a cell maintains a desired size for a given environment typically build on the deterministic setpoint perspective and focus on specific hypothesized paradigms, pithily described as sizer, timer, and adder (*15–20*). These approaches use a deterministic lens to build a framework that describes specific intragenerational phenomenology, and thence formulate a quasi-deterministic heuristic to motivate the requirement that cell size homeostasis be achieved intergenerationally (Figs. 1d, f). Data from related bacterial species analyzed in the Mother Machine occupy distinct positions along the continuum between specific paradigms (*21*), although the preponderance of attention has focused on the adder paradigm as applied to model bacterial systems (*22–25*). Theoretical extrapolations of intergenerational dynamics based on observed intragenerational phenomenology are often used to identify which of these schemes most accurately describes how growth control is actualized in different microorganisms (*18, 25, 26*).

**Fig. 1:**
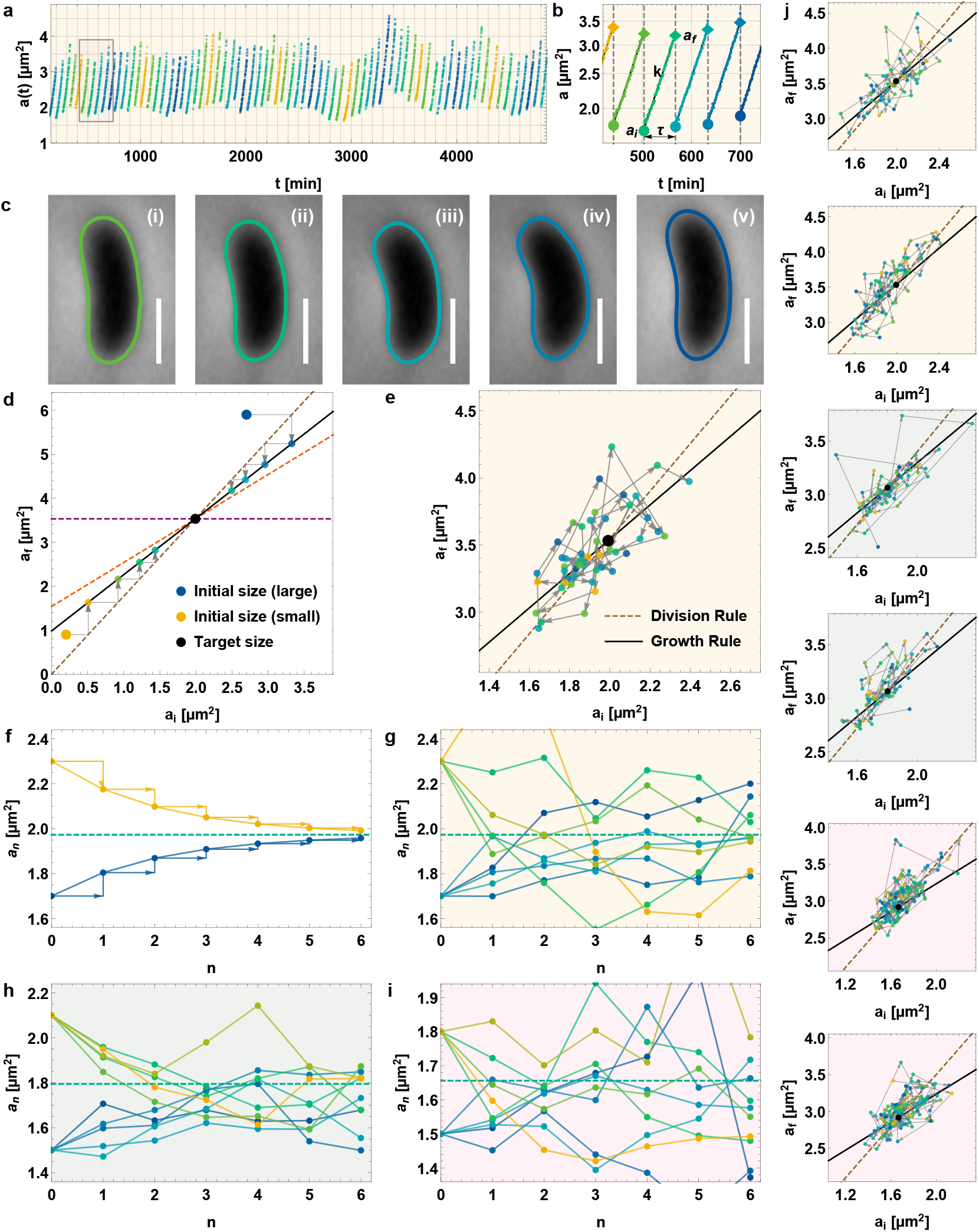
Stochastic intergenerational cell size trajectories of individual bacterial cells. (**a**)A typical individual *C. crescentus* stalked cell’s intergenerational growth and division trajectory obtained using our SChemostat technology under time-invariant balanced growth conditions. Colors indicate generations of the cell. (**b**) Exponential fits to growth trajectories of the generations corresponding to the box in (a), with initial size (*a*_*i,n*_, size at birth), final size (*a*_*f,n*_, size at division), growth rate (*k*), and interdivision time *(τ*) as indicated. The division ratio (*r*) relates *a*_*i,n*_ to *a*_*f,n*−1_. (**c**) Cell boundary splines generated by our custom automated image analysis routines, corresponding to *a*_*i,n*_ in the boxed region of (b). (**d**) Geometric interpretation of the heuristic argument for intergenerational homeostasis in the quasi-deterministic sizer–timer– adder paradigms. Dashed lines relate *a*_*f*_ to *a*_*i*_ in the sizer (*purple*, null slope), timer (*brown*, null intercept), and adder (*orange*, unit slope) mechanisms. The timer coincides with the “division rule”, which determines the size after division, *a*_*i,n*_ = *ra*_*f,n*−1_. The solid line is the experimentally measured “growth rule”, which allows the final size to be set by the initial size of the same generation. In this picture, cells deterministically adjust their sizes and exponentially relax to the target size set by the intersection between the growth and division rules. (**e**) Representative trajectory from our high-precision single-cell experimental observations showing stochastic intergenerational cell size homeostasis dynamics. Points correspond to single generations, and relate initial and final cell sizes (*a*_*i*_ and *a*_*f*_, respectively), within that generation. Successive generations are related by arrows. (**f**) Mean initial areas of successive generations that start from two different initial areas (1.6 and 2.0 *μ*m^2^, respectively). The dashed line represents the population mean of initial areas. (**g–i**) Initial areas from five randomly selected trajectories over successive generations for cells starting from the same initial areas as in (f), for cells growing in complex media (g), minimal media (h), and dilute complex media (i). These progressions are dramatically different from the prevailing quasi-deterministic sizer–timer–adder paradigm. Stochasticity is integral to intergenerational cell size homeostasis dynamics; there is no resemblance to the deterministic approach to a target size as suggested in (f). (**j**) Additional representative single-cell intergenerational cell size trajectories, plotted as in (e), for cells growing in complex media (*yellow background*), minimal media (*green background*), and dilute complex media (*pink background*), respectively. Each panel shows data from an individual cell.

However, as immediately evident in the dramatic contrast between the quasi-deterministic framework’s predictions (Fig. 1d) and high-precision, single-cell intergenerational data (Fig. 1e), these schemes may be unequivocally distinguished only in the deterministic case. In reality, homeostasis is an inherently stochastic process, whose study requires high-precision multigenerational datasets of statistically identical, non-interacting cells. A number of ingenious approaches have been developed to obtain single-cell datasets (*27–29*), but potential artifacts such as cell–cell contacts, mechanical confinement, and differential nutrient diffusion (*30*) reduce the precision in experimental control over growth conditions that each cell experiences (*31*). Because our SChemostat technology (*32, 33*) addresses each of these concerns, we used it to capture high-precision datasets of statistically identical, non-interacting *Caulobacter crescentus* cells over long periods of time (Fig. 1a–c), demonstrating that multigenerational trains of sizes of individual cells deviate dramatically from the naïve predictions derived from sizer–timer– adder frameworks (compare Fig. 1d to Figs. 1e, 1j, S1, S2, and S3; and Fig. 1f to Fig. 1g–i). Evidently, variations from the putative homeostatic setpoint are not only normal, but in fact an integral feature of the stochastic intergenerational homeostasis of an individual cell!

Taking a first principles-based approach deeply compatible with the underlying stochasticity, we previously observed remarkably precise intragenerational scaling laws (“emergent simplicities”) in single-cell growth between successive division events, including collapse of the mean-rescaled distributions of intragenerational cell size and of interdivision times from different conditions onto universal curves (*34*). However, as these intragenerational scaling laws do not guarantee intergenerational homeostasis, we sought to characterize the phenomenon of stochastic growth and division over many generations, without recourse to *a priori* assumptions about the machinery responsible for homeostasis.

## Direct evidence of stochastic intergenerational homeostasis of individual bacterial cell sizes

Surprisingly, direct evidence for stochastic intergenerational cell size homeostasis is not readily available in published works (*15–25*). Any attempt to confirm this hypothesis must meet both the necessary condition that mean cell size, averaged over an asynchronous population in balanced growth, be constant, as well as the sufficient condition that the distribution of cell sizes at birth (“initial size distribution”) be time-invariant. Here, we present individual-cell multigenerational SChemostat data from three different balanced growth conditions: cells grown in complex (undefined) media, minimal (defined) media, and dilute complex media. By experimentally confirming both the necessary and sufficient conditions (Figs. 2a–e, S4, and S5) we provide direct evidence of stochastic intergenerational homeostasis of individual cells.

**Fig. 2:**
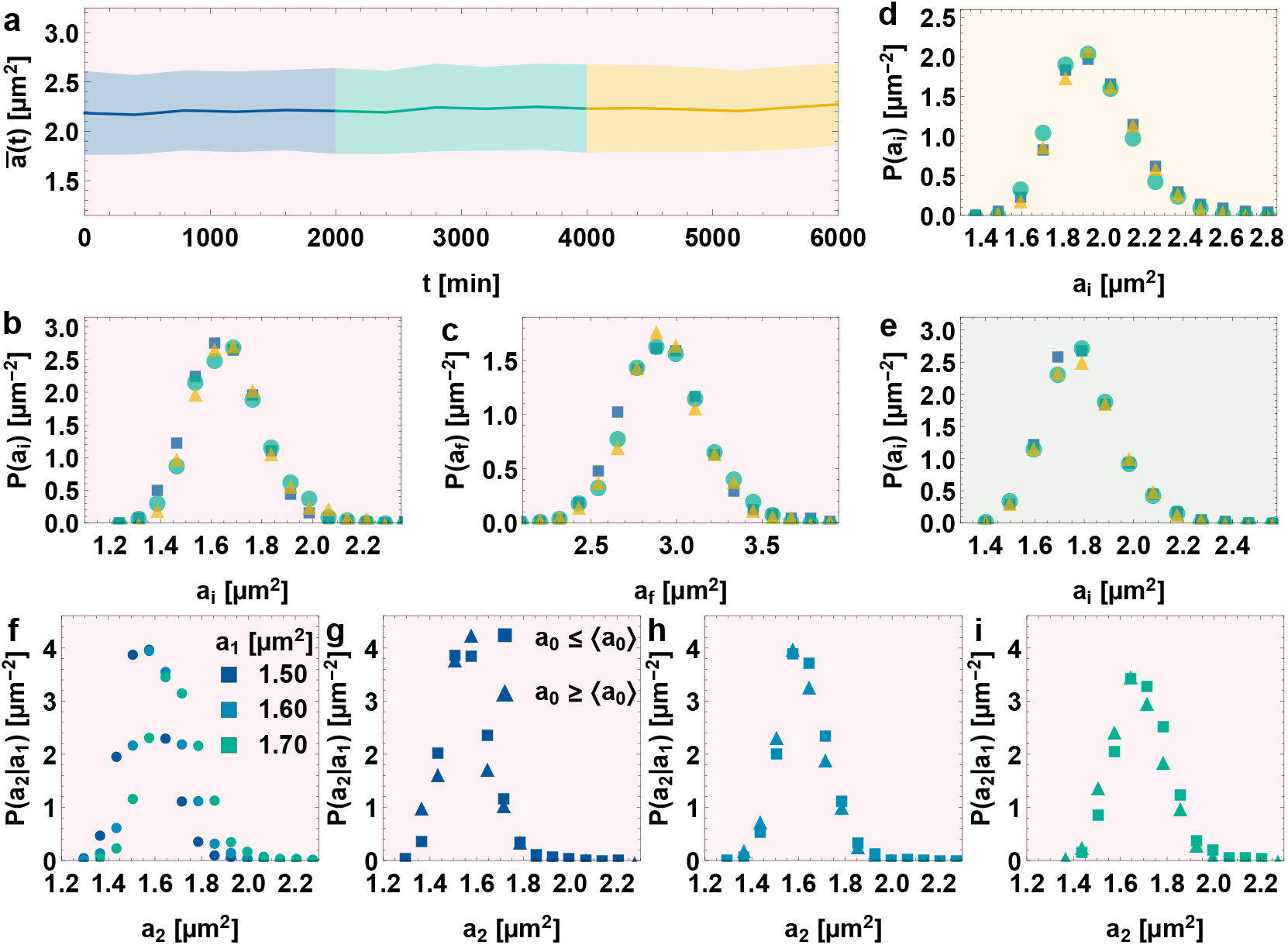
Precision measurements establish that individual cells maintain stochastic intergenerational cell size homeostasis. (**a**) The mean cell size averaged over an asynchronous population in balanced growth remains invariant when cell size homeostasis has been attained. Data are presented for cells growing in dilute complex media. (**b–c**) The distributions of pooled initial (b) and final (c) cell size are plotted for cell generations drawn from the three time windows shown in (a), using corresponding colors. These distributions are invariant, establishing that stochastic intergenerational homeostasis has been achieved and is sustained. (**d–e**) The distributions of pooled initial cell size are shown for cells growing in complex (d) and minimal (e) media, drawn from three windows as indicated by the point colors. (**f**) The distributions of the subsequent generation’s initial area (*a*_2_) given the current generation’s initial area (*a*_1_) are plotted for three different values of *a*_1_. Point colors indicate distinct values of *a*_1_. Data are taken from balanced growth conditions in dilute complex media. (**g–i**) For each value of *a*_1_ in (f), the conditional distribution of the subsequent generation’s initial area (*a*_2_) given the current generation’s initial area (*a*_1_) is further segmented according to whether the previous generation’s initial area (*a*_0_) was less than (*square*) or greater than (*triangle*) the population mean initial area. The distributions overlap irrespective of whether *a*_0_ was less than or greater than the population mean, indicating that *a*_2_ does not depend on *a*_0_ so long as the value *a*_1_ is fixed. Thus, initial size dynamics are Markovian.

## Emergent simplicity: Stochastic intergenerational cell size homeostasis is achieved through elastic adaptation

By analyzing successive values of initial cell sizes, we directly show that the intergenerational cell size dynamics are Markovian. Briefly, given a specified generation’s initial size (*a*_1_), the distribution of the next generation’s initial size (*a*_2_) is completely determined and does not depend on the preceding generations’ initial sizes (*a*_0_, *a*_−1_, and so on) (Figs. 2f–i, S6, and S7, and Supplementary Materials (SM) Mathematical Derivations Sec. 1). In other words, *P*(*a*_*n*_|*a*_*n*−1_, *a*_*n*−2_, *a*_*n*−3_ …) = *P*(*a*_*n*_|*a*_*n*−1_) = *P*_1_(*a*_*n*_|*a*_*n*−1_) for any n under balanced growth conditions. This result is key to building the intergenerational theoretical framework. The immediate implication is that, under balanced growth conditions, intergenerational cell size homeostasis is maintained through reflexive, elastic adaptation.

## Theoretical formulation of stochastic intergenerational framework for cell size homeostasis

Using the key emergent simplicity that the intergenerational cell size dynamics are Markovian, the basic equation governing the multigenerational dynamics of initial cell sizes (SM Mathematical Derivations Sec. 2) is:

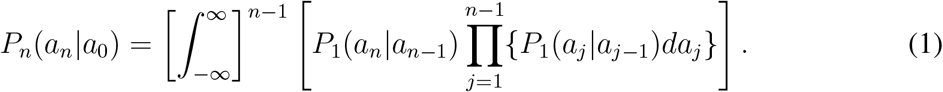

In this formulation, homeostasis is achieved if *P*_*n*_(*a*_*n*_|*a*_0_) converges to the same well-behaved distribution for all values of initial cell sizes, *a*_0_, after a sufficient number of generations (*n* ≫ 1) have elapsed. This is indeed true for our data in all three conditions (Figs. 3e, f, and k–n).

**Fig. 3:**
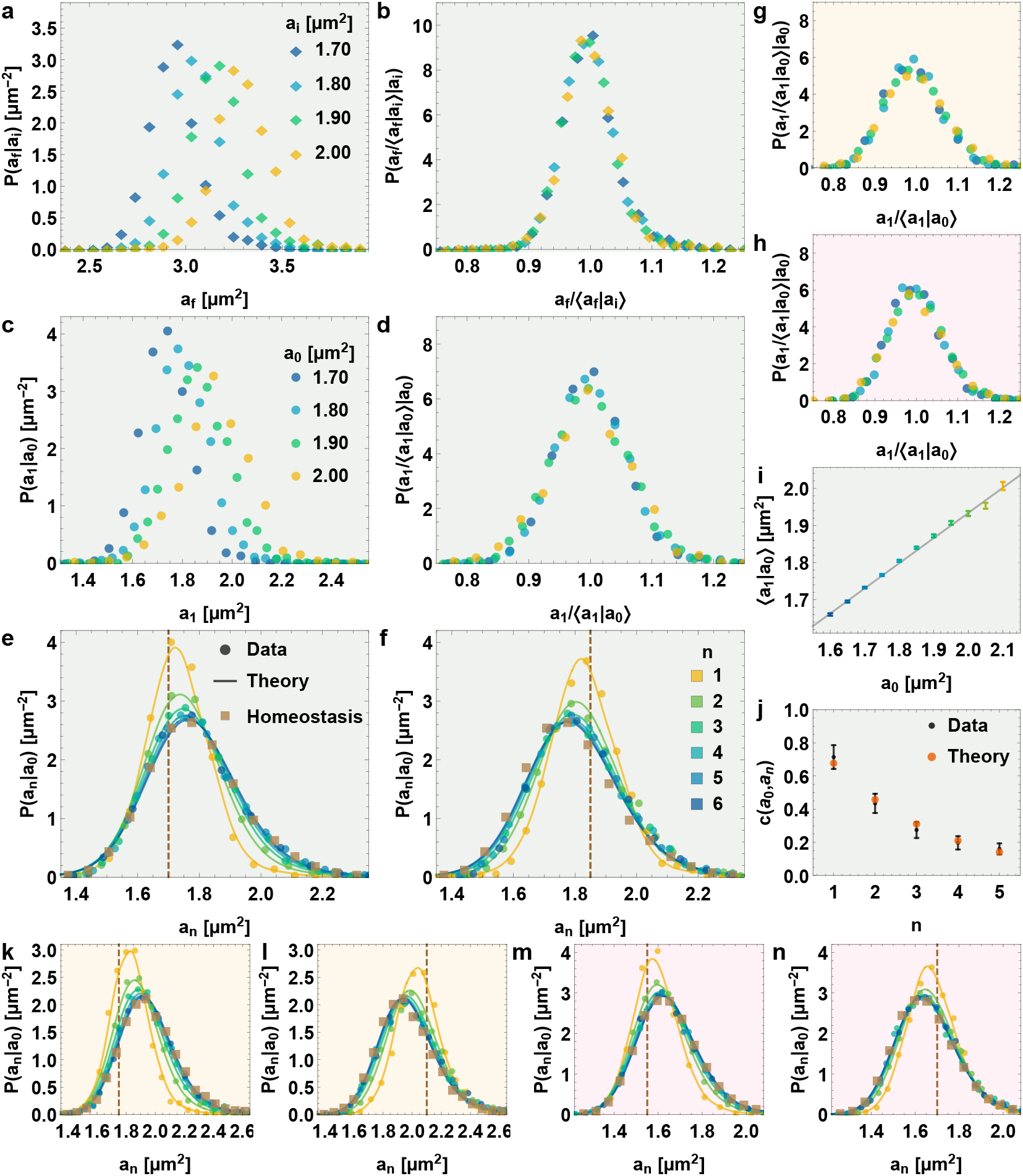
Emergent simplicity: intergenerational scaling law governing stochastic individual cell size homeostasis. (**a**) *P*_*f*_ (a_f_ | a_i_), the distribution of final size, *a*_*f*_, pooled by distinct values of initial size, *a*_*i*_, within the same generation. Data are shown for cells growing in minimal media. Point colors indicate distinct values of *a*_*i*_. (**b**) Corresponding mean-rescaled distributions, with point colors as indicated in (a); ⟨*a*_*f*_ |*a*_*i*_ = *μ*_*f*_ (*a*_*i*_) is the mean *a*_*f*_ for a given *a*_*i*_. (**c**) *P*(*a*_1_ | *a*_0_), the distribution of the subsequent generation’s initial size, *a*_1_, pooled by distinct values of the current generation’s initial size, *a*_0_, for cells growing in minimal media. Point colors indicate distinct values of *a*_0_. (**d**) Mean-rescaled distributions corresponding to (c). Thus, for a given growth condition, the two-dimensional function *P*(*a*_1_ | *a*_0_) (c) is constructed from two one-dimensional functions: the mean-rescaled distribution (d) and the *a*_0_-dependent mean ⟨*a*_1_| *a*_0_ ⟨= *μ*(*a*_0_) shown in (i), which sets the calibration curve for how to “stretch” the *a*_0_-independent rescaled distributions in (d) to recover the a_0_-dependent distributions in (c). (**e–f**) Intergenerational evolution of the distribution of initial cell sizes, *a*_*i*_, beginning from a given initial size in the starting generation (location of dashed line), for two different starting values of *a*_*i*_. Data are shown for cells growing in minimal media. *n* denotes the generation number, with the starting generation labeled by *n* = 0. Points correspond to experimental data and curves to fitting-free predictions from our theoretical intergenerational framework, which takes as the only inputs the directly measured mean-rescaled distribution (d) and mean calibration curve (i). The compelling fitting-free data–theory match independently validates the emergent simplicities used as inputs for our theoretical framework: the Markovian nature of intergenerational cell size dynamics and *a*_*i*_-independence of the mean-rescaled size distribution. (**g–h**) Mean-rescaled distributions as in (d), but for cells growing in complex (g) and dilute complex (h) media. (**i**) The binned means of the next generation’s initial areas plotted for a range of the current generation’s initial areas, along with the linear fit. This calibration curve is used to restore the full conditional next generation’s initial size distributions conditioned on the current generation’s initial size (c) from the corresponding mean-rescaled distributions (d). (**j**) The coefficient of correlation between the initial areas in the initial (zeroth) generation and after n consecutive generations for different n. The correlations predicted by our fitting parameter-free theoretical framework (*orange*) closely match those measured from data (*black*). (**k–n**) Distributions corresponding to (e–f) for cells growing in complex (k–l) and dilute complex (m–n) media, respectively.

## Emergent simplicity: A scaling law governing stochastic intergenerational homeostasis

We then extracted from our data the conditional probability distribution of the final size conditioned on the initial size within the same generation, *P*_*f*_ (*a*_*f*_ |*a*_*i*_), and found a remarkable emergent simplicity (Figs. 3a, 3b, S8): the mean-rescaled distribution is independent of the value of the initial size. Thus,

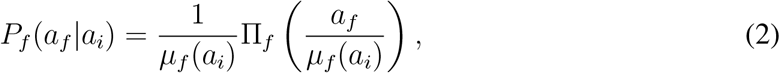

where П_*f*_ is the mean-rescaled distribution with unit mean and *μ*_*f*_ (*a*_*i*_) = ⟨ *a*_*f*_ |*a*_*i*_⟩ is the mean of the final area, *a*_*f*_, as a function of the initial area, *a*_*i*_. This observation holds for all three conditions. This emergent simplicity facilitates our task of developing the theory of stochastic intergenerational homeostasis; instead of considering *P*_*f*_ (*a*_*f*_ |*a*_*i*_), which is generally a function of two variables, we need only work with two one-dimensional functions, П_*f*_ and *μ*_*f*_ . Of these, *μ*_*f*_ (*a*_*i*_) simply serves as a “calibration curve” for restoring the unscaled distribution from the universal mean-rescaled distribution for a given growth condition. From data, *μ*_*f*_ (*a*_*i*_) is well approximated by a straight line (Fig. S10).

## The exact stochastic map governing intergenerational cell size dynamics and homeostasis is directly validated by fitting-free data–theory comparisons

Taken together, these two emergent simplicities—the Markovian nature of intergenerational cell size dynamics (Figs. 2f–i, S6, and S7) and the intergenerational scaling law (Eq. 2, Figs. 3a, 3b, S8)—serve as starting points to motivate and develop a data-informed theoretical framework. Furthermore, these data reveal yet another emergent simplicity: the division ratio, r, defined as the ratio of the initial size in a given generation to the final size in the previous generation, is independent of the final size in the previous generation. Thus, *P*(*r*|*a*_*f*_) = *P*(*r*) is independent of *a*_*f*_; when combined with Eq. (2), this implies that the conditional distribution of the next generation’s initial size, conditioned on the current generation’s initial size, must also undergo a scaling collapse upon rescaling by the conditional mean (Figs. 3c, 3d, 3g, 3h, S9):

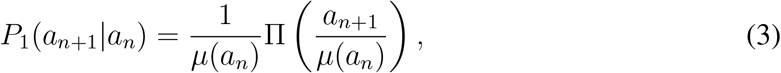

where П denotes the new mean-rescaled distribution (with unit mean) and *μ*(*a*_*n*_) = ⟨*a*_*n*+1_|*a*_*n*_⟩ the conditional mean of the initial size in the next generation, given the initial size in the current generation. These functions can be related to the corresponding quantities in Eq. (2): П(*s*) = ⟨*r*^−1^П_*f*_ (*s* ⟨*r*⟩ /*r*) ⟩_*r*_ ⟨r⟩ and *μ*(*a*) = ⟨*r*⟩ *μ*_*f*_ (*a*), where ⟨*r*⟩ is the mean division ratio and ⟨· ⟩_*r*_ denotes averaging over fluctuations of r (SM Mathematical Derivations Sec. 3). We see direct evidence of this intergenerational scaling of initial sizes for all three conditions (Fig. S9).

Using this intergenerational scaling law and the Markovian framework in Eq. (1), we can predict the complete stochastic map for intergenerational cell size statistics of an individual cell in a given balanced growth condition by iterating the Langevin-type map *a*_*n*+1_ = *s*_*n*_*μ*(*a*_*n*_),where (.. ., *a*_*n*−1_, *a*_*n*_, *a*_*n*+1_,…) are random variables corresponding to possible trajectories of cell size, and the random variable *s*_*n*_ is drawn from the distribution П in Eq. (3), independently for each *n*. The experimental inputs are the universal mean-rescaled distribution function, П, and the calibration curve corresponding to the conditional mean, *μ*, for that growth condition (SM Mathematical Derivations Sec. 3). Our fitting-free predictions match our data remarkably well for all conditions (Figs. 3e, f, and k–n). Thus, we have written down a data-validated fitting-free theoretical framework: the stochastic map governing stochastic intergenerational homeostasis!

## Universality of the stochastic map governing stochastic intergenerational homeostasis

The results presented here are broadly valid across different growth conditions, morphologies, experimental modalities, and organisms. We have found that they hold for cell growth in different temperatures spanning the entire physiological range of *C crescentus*, as well as for a rod-shaped mutant of *C. crescentus*, which follows the same intergenerational scaling law, Eq. (3), and hence is governed by the same stochastic map described here. These data were collected using the SChemostat, which ensures statistically independent, non-interacting cells; however, because its application is limited to microbes that can be made conditionally adhesive, the Mother Machine remains the go-to device for a wide range of microbes. To benchmark data obtained from Mother Machine-based technologies, and to assess the effects from unwanted factors such as mechanical stress, in (*31*) we developed a protocol to study comparable data from rod-shaped *C. crescentus* in both devices to obtain the first heads-on comparison of high-precision data of growth and division dynamics of single bacterial cells from different experimental apparatuses. Despite clear differences in observable growth and division data from the SChemostat and Mother Machine, we established a principled route to account for the impacts of specific experimental modalities on the observed single-cells dynamics and established that the cells in both setups followed the same intergenerational scaling law and cell size dynamics as reported here! We have also reanalyzed published data for *Escherichia coli* and *Bacillus subtilis* obtained using the Mother Machine (*17*), alongside our own data for *C. crescentus* (*31, 35*), and again find that the intergenerational scaling law reported here also governs their stochastic cell size homeostasis.

## Stochasticity specifies the conditions for homeostasis

In addition to fully determining the stochastic map governing intergenerational homeostasis, the observed intergenerational scaling law also determines the general necessary and sufficient conditions for stochastic homeostasis: given the observed linear form of the conditional mean, *μ*(*a*) = *αa* + *β*, the magnitude of the slope, *α*, must be less than or equal to 1/*s*_max_, where *s*_max_ is the maximum allowed value of the variable governed by the mean-rescaled probability distribution П in Eq. (3) (see (*35*) for details of derivation). Conversely, given *α*, the upper bound of the П distribution must lie below 1/*α*. This prediction is indeed consistent with our data; the values of 1/*α* corresponding to Figs. 3 g, d, and h (equivalently, Figs. S9 b, d, and f) are ∼ 1.6, 1.5, and 1.7 (obtained from Figs. S10 b, e, and h), respectively. In each case, the mean-rescaled distribution is evidently bounded from above by the corresponding value of 1/*α*. Since П has unit mean, the presence of stochasticity implies that *s*_max_ > 1; thus, our homeostasis requirement, |*α*| ≤ 1/*s*_max_, is a stronger bound on *α* than that obtained from the quasi-deterministic sizer–timer–adder paradigm of |*α*| < 1 (*22, 36–38*). Violation of the condi-tion leads to breakdown of homeostasis, manifested as appearance of heavy tails and divergent higher-order moments of the long-time size distribution (*35*). As *α* > 1/*s*_max_ increases towards 1, i.e., the violation of the upper bound becomes more pronounced, more moments (especially those of lower order) diverge until all semblance of the homeostatic size distribution is lost when *α* crosses to values over 1.

## Reconciling the mythical “average” cell with the representative stochastic cell

Combining our intergenerational framework with the linear nature of the conditional mean, we find that when the starting cell size is larger or smaller than the homeostatic mean size, the mean size in subsequent generations relaxes exponentially to the homeostatic target, with an exponential rate given by ln(1/|*α*|) (*35*). Consequently, the correlation function *C*_*mn*_ of cell sizes in generations *m* and *n* also decays exponentially with rate ln(1/|*α*|) (SM Mathematical Derivations Sec. 4):

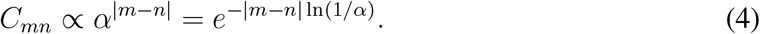

Figs. 3j and S10c, f, and i show good agreement between this theoretical prediction and the experimentally calculated correlation functions. This mean-level behavior is reminiscent of the quasi-deterministic evolution of cell size with generation, which is predicted to exponentially relax to the setpoint size with the same exponential rate, ln(1/|*α*|). Thus, the quasi-deterministic picture describes an *average* cell, but not a *representative* cell (*39*).

## Concluding remarks

Our approach of identifying “top down” organism-level emergent simplicities complements the common atomistic view that understanding essentially complex phenomena in essentially complex systems, such as whole organisms, must focus on the dissection of genetic and signaling networks (*40*). Yet the identification of these emergent simplicities, or “constraints that deconstrain” (*41, 42*), can serve to provide non-trivial insights into the underlying mechanism. Through purely theoretical considerations, in (*43*) we have identified architectural underpinnings of the intergenerational scaling law reported here. The intergenerational scaling law, Eq. (3), arises naturally when the commitment to the division process is triggered by the thresholding of a biochemical species whose copy numbers increase exponentially in proportion to cell size, following which the cell divides after a stochastic time delay. The details of the implementation of the mechanism, such as the molecular identity of the biochemical species, the specific implementation in a given organism or growth condition, and other model parameters, can vary. In other words, these details are “deconstrained”. Specifically, in *C crescentus*, these dynamics qualitatively match the known behavior of FtsZ, which is responsible for the formation of the Z-ring leading to the triggering of cell constriction. Yet, generally, we expect that the control system governing stochastic size homeostasis shares the core features we have identified, leading naturally to the emergent simplicities reported here.

In sum, our results transcend the minutiae of the condition-specific molecular networks implicated in growth control in distinct environmental conditions and organisms. While the intergenerational scaling law holds across multiple balanced growth conditions and bacterial species, the shape of the collapsed distribution may change from condition to condition. The mechanism is universal, but the specific instantiation depends on the characteristics of the balanced growth condition. Intriguingly, the data presented here show early indications of a “precision–speed” tradeoff (*44*), encapsulated in the different shapes of the mean-rescaled curves from different growth conditions—narrower for the slower growth condition (defined minimal media) and broader for the faster growth condition (undefined complex media). An analogous precision– energy tradeoff is indicated in the observation that the slope, *α*, of the mean calibration curve, *μ*(*a*_*i*_) = *αa*_*i*_ + *β*, is comparatively closer to the *upper* limit 1/*s*_max_ set by our intergenerational framework, and not a lower value such as 0 that would correspond to perfect size control. What costs and benefits would determine the tradeoffs and optima in these contexts? These promising lines of future inquiry will build on the results presented here.

Finally, despite the long-standing criticism directed at the simplicity of the memory-free design of Ashby’s Homeostat, our work shows it to be a representative model for natural homeostasis in a living organism, after all! Perhaps this is the most startling and elegant simplicity uncovered: over intergenerational timescales, cell size homeostasis is an elastically adapting process. While excitement has been building around the general possibility that single cells may serve as useful arenas for exploring the phenomenology of learning without neurons (*45, 46*), our work shows that the intergenerational evolution of bacterial cell size, under balanced growth conditions, is history-independent and therefore cannot serve as a diffuse living brain with a capacity for learning on intergenerational timescales. Where then is a bacterial cell’s intergenerational memory manifested? Early indications are that the instantaneous single-cell growth rate may be the answer (*47, 48*).

## Acknowledgments

S.E. and S.I.-B thank Purdue’s Data Mine Learning Community for computational resources. We are grateful to the Iyer-Biswas group members for insightful discussions and feedback. S.I.-B. thanks Daniel S. Fisher for useful questions.

## Author contributions

K.J. and S.I.-B. conceived of the research; K.J., R.R.B. and S.I.-B. designed the research; K.J. extracted the intergenerational scaling law from the data and performed data analysis under the guidance of S.I.-B.; K.J., R.R.B. and S.I.-B. developed the theoretical framework and performed analytic calculations; C.S.W. and S.I-.B. designed the experimental setup; C.S.W. and K.F.Z. actualized and refined the experimental pipeline under the guidance of S.I.-B.; C.S.W. and S.E. designed and developed the analysis pipeline with input from R.R.B. and S.I.-B.; C.S.W, K.F.Z., E.S., J.C. and S.I.-B. performed experiments; K.J., C.S.W., R.R.B. and S.I.-B. wrote the paper with input from all authors; S.I.-B. supervised the research.

## References

1. G. E. Billman, Frontiers in Physiology 11 (2020).

2. C. Bernard, Introduction á l’eétude de la meédecine expeérimentale, no. 2 (JB Baillière, 1865).

3. W. B. Cannon, Á Charles Richet: ses amis, ses collègues, ses eélèves (in French), A. Pettit, ed. (Les EÉ ditions Meédicales, Paris, 1926), p. 91.

4. W. B. Cannon, The Wisdom of the Body (W.W. Norton & Company, Inc., New York, 1932).

5. J. D. Hardy, Harvey Lectures 49, 242 (1953–1954).

6. N. Wiener, Cybernetics: or Control and Communication in the Animal and the Machine (MIT Press, Cambridge, MA, 1948).

7. M. E. Kotas, R. Medzhitov, Cell 160, 816 (2015).

8. S. Mukherjee, The Song of the Cell: An Exploration of Medicine and the New Human (Simon and Schuster, 2022).

9. S. Iyer-Biswas, F. Hayot, C. Jayaprakash, Phys. Rev. E 79, 031911 (2009).

10. S. Iyer-Biswas, Applications of methods of non-equilibrium statistical physics to models of stochastic gene expression, Ph.D. thesis, Ohio State University (2009).

11. S. Iyer-Biswas, A. Zilman, First-Passage Processes in Cellular Biology (John Wiley & Sons, Inc, 2016), chap. 5, pp. 261–306.

12. F. Jafarpour, M. Vennettilli, S. Iyer-Biswas, arXiv:1703.10058 (2017).

13. W. R. Ashby, Radio Electronics pp. 77–79 (March Issue, 1949).

14. Life Magazine 33, 45 (1952).

15. T. W. Spiesser, C. Muüller, G. Schreiber, M. Krantz, E. Klipp, The FEBS Journal 279, 4213 (2012).

16. M. Deforet, D. van Ditmarsch, J. B. Xavier, Biophysical Journal 109, 521 (2015).

17. S. Taheri-Araghi, et al., Current Biology 25, 385 (2015).

18. J. T. Sauls, D. Li, S. Jun, Current Opinion in Cell Biology 38, 38 (2016).

19. M. M. Logsdon, et al., Current Biology 27, 3367 (2017).

20. L. Willis, K. C. Huang, Nature Reviews Microbiology 15, 606 (2017).

21. S. Jun, S. Taheri-Araghi, Trends in Microbiology 23, 4 (2015).

22. A. Amir, Physical Review Letters 112, 208102 (2014).

23. M. Campos, et al., Cell 159, 1433 (2014).

24. J. Lin, A. Amir, Cell Systems 5, 358 (2017).

25. F. Si, et al., Current Biology 29, 1760 (2019).

26. K. R. Ghusinga, C. A. Vargas-Garcia, A. Singh, Scientific Reports 6, 30229 (2016).

27. P. Wang, et al., Current Biology 20, 1099 (2010).

28. J. R. Moffitt, J. B. Lee, P. Cluzel, Lab on a Chip 12, 1487 (2012).

29. T. M. Norman, N. D. Lord, J. Paulsson, R. Losick, Nature 503, 481 (2013).

30. L. Potvin-Trottier, S. Luro, J. Paulsson, Current Opinion in Microbiology 43, 186 (2018).

31. K. F. Ziegler, et al., Molecular Biology of the Cell 35:ar78, 1 (2024).

32. S. Iyer-Biswas, et al., Proceedings of the National Academy of Sciences of the United States of America 111, 15912 (2014).

33. F. Jafarpour, et al., Physical Review X 8, 021007 (2018).

34. S. Iyer-Biswas, G. E. Crooks, N. F. Scherer, A. R. Dinner, Physical Review Letters 113 (2014).

35. K. Joshi, R. R. Biswas, S. Iyer-Biswas, bioRxiv:2023.01.20.525000 (2023).

36. L. Susman, et al., Proceedings of the National Academy of Sciences of the United States of America 115, E5679 (2018).

37. W. F. Marshall, Annual Review of Biophysics 45, 49 (2016).

38. M. B. Ginzberg, R. Kafri, M. Kirschner, Science 348, 1245075 (2015).

39. S. Sanders, K. Joshi, P. A. Levin, S. Iyer-Biswas, PLoS Genetics 19, e1010505 (2023).

40. L. H. Hartwell, J. J. Hopfield, S. Leibler, A. W. Murray, Nature 402, C47 (1999).

41. M. Kirschner, J. Gerhart, Proc. Natl. Acad. Sci. U. S. A. 95, 8420 (1998).

42. J. C. Doyle, M. Csete, Proc. Natl. Acad. Sci. U. S. A. 108 Suppl 3, 15624 (2011).

43. K. Joshi, C. S. Wright, R. R. Biswas, S. Iyer-Biswas, bioRxiv:2023.11.15.567256 (2023).

44. G. Lan, P. Sartori, S. Neumann, V. Sourjik, Y. Tu, Nature Physics 8, 422 (2012).

45. J. Gunawardena, Proceedings of the IEEE 110, 590 (2022).

46. D. Rajan, et al., Current Biology 33, 241 (2023).

47. K. Joshi, et al., bioRxiv:2023.05.27.542601 (2023).

48. K. Joshi, S. Roy, R. R. Biswas, S. Iyer-Biswas, bioRxiv:10.1101/2023.03.07.531540 (2023).

49. C. S. Wright, et al., Scientific Reports 5, 9155 (2015).

